# *In vitro* characterization of the yeast DcpS, a scavenger mRNA decapping enzyme

**DOI:** 10.1101/644492

**Authors:** Madalee G. Wulf, John Buswell, Siu-Hong Chan, Nan Dai, Katherine Marks, Evan R. Martin, George Tzertzinis, Joseph M. Whipple, Ivan R. Corrêa, Ira Schildkraut

## Abstract

Eukaryotic mRNAs are modified at their 5’ end early during transcription by the addition of *N*7-methylguanosine (m^7^G), which forms the “cap” on the first 5’ nucleotide. Identification of the 5’ nucleotide on mRNA is necessary for determination of the Transcription Start Site (TSS). We explored the effect of various reaction conditions on the activity of the yeast scavenger mRNA decapping enzyme DcpS (yDcpS) and examined decapping of 30 chemically distinct cap structures varying the state of methylation, sugar, phosphate linkage, and base composition on 25mer RNA oligonucleotides. Contrary to the generally accepted belief that DcpS enzymes only decap short oligonucleotides, we found that yDcpS efficiently decaps RNA transcripts as long as 1400 nucleotides. Further, we validated the application of yDcpS for enriching capped RNA using a strategy of specifically tagging the 5’ end of capped RNA by first decapping and then recapping it with an affinity-tagged guanosine nucleotide.

## Introduction

The 5’ ends of eukaryotic messenger RNA (mRNA) carry a 5’ to 5’ triphosphate linked 7-methyl-guanosine (m^7^G) cap. The capping occurs at an early stage of transcription and is involved in splicing, stabilizing and directing the transcript to the cytoplasm where it is translated. In higher eukaryotes, the canonical m^7^G cap can be further modified by 2’-*O*-methylation at nucleotide positions +1 (Cap 1) and +2 (Cap 2) of the RNA. Both capping and decapping of mRNA are critical components of gene expression. The enzymes responsible for the decapping have been described from two distinct protein families, the Nudix pyrophosphohydrolases^1^ and the HIT family of pyrophosphatases^2,3^. The distinguishing characteristic of catalysis by the two families of decapping enzyme is the resulting products of the reactions (Figure 1). Nudix decapping enzymes perform metal dependent hydrolysis of the phosphodiester bond between the beta and alpha phosphates to leave a 5’-monophosphate RNA (*p*RNA) and a 7-methyl-guanosine diphosphate (m^7^GDP). A few exceptions have been reported where a Nudix decapping activity results in mixed diphosphate and monophosphate 5’ RNA ends^1^. On the other hand, HIT decapping enzymes, including the so-called scavenger decapping enzyme DcpS, hydrolyze the phosphodiester bond between the gamma and beta phosphates to leave a 5’-diphosphate-RNA (*pp*RNA) and a 7-methyl-guanosine monophosphate^4,5^ (m^7^GMP) and are independent of divalent metal ions^6^.

**Figure 1.**
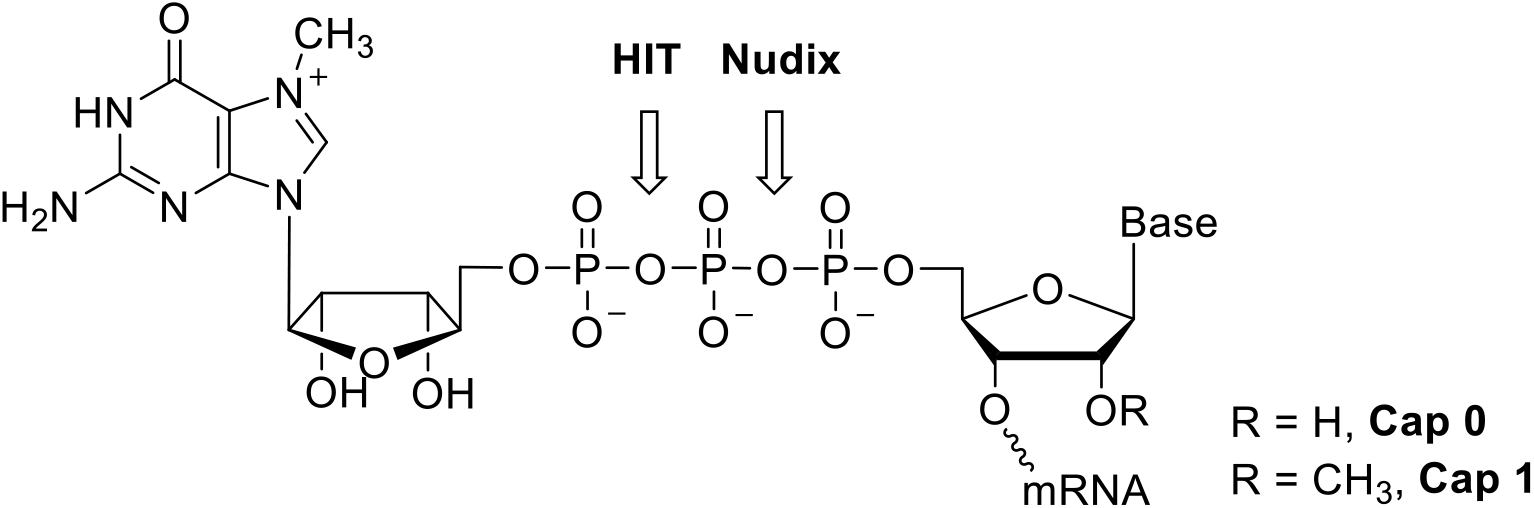
Cap structure and site of hydrolysis for the HIT and Nudix enzymes.

DcpS proteins of the HIT family have been identified in fungi and animals. DcpS contains a conserved set of histidines, the histidine triad sequence, which is necessary for catalysis. These enzymes are reported to “scavenge” the m^7^GMP from the 5’ terminus of short mRNA fragments that are produced after extensive 3’ to 5’ exonuclease digestion by the exosome complex. Capped RNAs of a length of ten nucleotides and longer have been reported to be resistant to decapping with the human DcpS^4,7^. Both human and yeast DcpS have been shown to play a biological role in the 3’ to 5’ mRNA decay pathway^4,8^.

Here we characterize the yeast, *Saccharomyces cerevisiae*, DcpS (yDcpS) with respect to cap structure specificity and RNA substrate length. The focus of our study is not to investigate the biological role of DcpS, but rather to study the enzyme *in vitro* in order to establish conditions in which it might be useful for manipulating RNA. We demonstrate that yDcpS under specified conditions can decap RNA of a length up to at least 1.4 kb, thus rendering the 5’ end of the RNA enzymatically “recappable”. In addition, we have determined the *in vitro* specificity of yDcpS with respect to 30 different cap analog structures at the 5’ end of a 25mer RNA substrate. These analogs include Cap 0 and Cap 1 structures varying the nucleotide at position +1 of the RNA (A, m^6^A, C, G, or U), as well as the identity (G, dG, araG, I, A, C, U, or nicotinamide) and methylation status (m^7^G, m^2,2,7^G and unmethylated G) of the cap nucleotide, and the length of the internucleotidic phosphate bridge (di-, tri- or tetraphosphate). Our results indicate that DcpS can decap most guanosine caps; in contrast, it shows no activity towards adenosine, cytidine or uridine capped RNAs.

In a previous work, we developed a method termed Cappable-seq^9^ to enrich primary prokaryotic RNA transcripts by capping their 5’ triphosphate with 3’-desthiobiotin-GTP (DTB-GTP). In an effort to extend this method to eukaryotic mRNA, we demonstrate here that yDcpS can decap the 5’ end of m^7^G-capped RNA transcripts of a length from 90 to 1400 nucleotides without appreciable length bias. Further, the yDcpS-treated transcripts can be recapped with DTB-GTP and recovered after binding to, and eluting from, streptavidin beads.

## Results

Liu *et al.*^4^ and Wypijewska *et al.*^10^ presented data indicating that human DcpS (hDcpS) does not decap RNAs longer than approximately ten nucleotides. Cohen *et al.* showed that the nematode DcpS was even more limited in its ability to decap oligonucleotides longer than 3 nucleotides^11^. This precept of acting solely on capped short oligonucleotides has been attributed to all homologous HIT enzymes^12^, despite Salehi *et al.*^13^ identifying a HIT protein in *Schizosaccharomyces pombe*, Nhm1, which was determined to decap long mRNA. Nhm1 has about 70% amino acid sequence similarity to the yeast and human DcpS enzymes. As these two views of Milac and Salehi are conflicting, we decided to characterize yDcpS with respect to its ability to decap RNA of different lengths. We used recombinant yDcpS expressed in *E. coli* and purified to homogeneity as described in Methods.

### Pyrophosphorolysis of the m^7^GpppA dinucleotide

We first looked at catalysis of the dinucleotide cap analog m^7^GpppA, as the characterization of DcpS typically involves pyrophosphorolysis of cap analogs which are dinucleotides of NpppN structure^4,6,14–16^. Cap analogs can be considered as a capped RNA of one nucleotide in length. Both Malys *et al.*^17^ and Liu *et al.*^18^ demonstrated the pyrophosphorolysis of m^7^GpppG dinucleotide with a rate constant of 0.012/sec for yDcpS and 0.09/sec for hDcpS, respectively. We compared the relative activity of hDcpS and yDcpS towards the m^7^GpppA dinucleotide. We determined the decapping rate at 50 μM m^7^GpppA dinucleotide by LC/MS analysis (Figure 2A). The results indicate the two enzymes catalyze the pyrophosphorolysis of dinucleotide with similar rates: yDcpS at 0.08/sec and hDcpS at 0.14/sec. Although the rate constant seen here for yDcpS is about 7 times faster than that reported by Malys *et al.*, their incubation temperature was lower (30 °C), the ionic strength was greater (100 mM KOAc) and the pH was higher (pH 7.0). Thus, the catalysis of the dinucleotide observed with our preparation of yDcpS is consistent with published literature.

**Figure 2.**
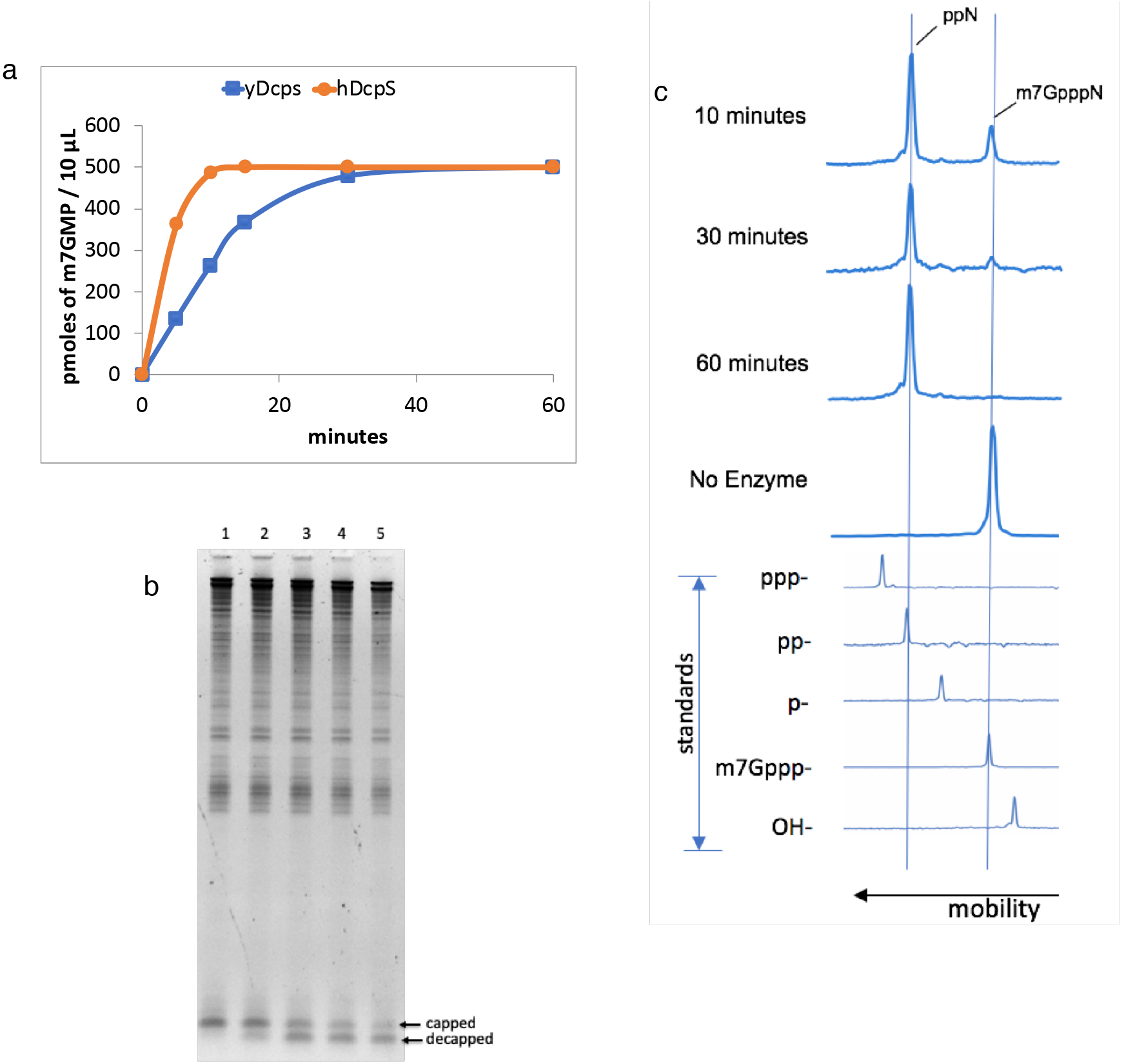
Biochemical Assays for DcpS. (**a**) Conversion of m^7^GpppA dinucleotide to m^7^GMP. A 100 μL reaction in decapping buffer containing 50 μM m^7^GpppA and either 50 nMy DcpS (blue □) or 85 nM hDcpS (red ○) was incubated at 37 °C. 10 μL samples were withdrawn at the indicated times. The reactions were terminated by the addition of equal volume of phenol/chloroform. The relative quantity of the remaining dinucleotide m^7^GpppA, m^7^GMP, and ADP were determined by LC-MS. (**b**) Decapping of a 25mer Cap 1 RNA by yDcpS. A 30 μL reaction containing 300 ng of total *E. coli* RNA and 60 ng of 25mer Cap 1 RNA was incubated for 3 hours at 37 °C with 130 ng of yDcpS in 10 mM MES pH 6.5 and 1 mM EDTA. At 0 minutes a 5 μL aliquot was mixed with 2X RNA loading dye stop solution (Lane 1). Likewise 5 μL aliquots were taken at 5, 60, 120, and 180 min (Lanes 2 to 5, respectively). The RNA aliquots were analyzed by 15% TBE-Urea PAGE stained with SYBR Gold. (**c**) Capillary electrophoresis of 3’-FAM-labeled 5’-capped 25mer RNA. Top panel shows a representative example of the decapping reaction progress. The 25mer Cap 0 RNA was incubated with yDcpS for various times indicated on the left. Bottom panel shows the mobility shift for standards of 25mer RNAs containing different 5’ ends. All data were plotted relative to GeneScan™120 LIZ™ Applied Biosystems standards.

### Yeast DcpS decaps a 25mer RNA

To determine whether yDcpS can decap an RNA longer than 15 nucleotides, a 25mer Cap 1 RNA synthesized and capped *in vitro*, was combined with total *E. coli* RNA and used as a substrate. The *E. coli* RNA was included to more closely resemble an *in vitro* reaction where only a subset of total eukaryotic RNA would be substrate for the DcpS. This mixture was incubated with yDcpS, and aliquots were sampled over time and analyzed by gel electrophoresis. As shown in Figure 2b, with increasing incubation time of up to three hours, a larger fraction of the 25mer RNA is shifted to the position of the decapped species, demonstrating that yDcpS is capable of decapping a longer RNA than what was reported previously.

### The effect of ionic strength and pH on decapping activity

*In vitro* decapping of DcpS was assessed at various ionic strength and pH conditions by using a synthetic 25mer Cap 0 RNA modified with a 3’-FAM label. Substrate and product of the decapping reaction were resolved and quantified by capillary electrophoresis (CE) (Figure 2c). As shown in Table 1A, the decapping reaction is significantly inhibited by increasing salt concentration, with KCl being a more potent inhibitor than NaCl. The optimal buffering pH was in the 6 - 6.5 range. yDcpS exhibited very little decapping activity in the VCE Capping Buffer at pH 8.0 and in phosphate buffer at pH 7.0 (Table 1B). Furthermore, a control reaction with a 5’ triphosphate 3’-FAM labeled 25mer showed no measurable conversion of the triphosphate to di- or mono-phosphate after incubation with 50 μM yDcpS in decapping buffer at 37°C for one hour as determined by CE analysis.

**Table 1.** Summary of the effects of reaction conditions on the yDcpS activity. *In vitro* decapping of 5’-capped synthetic 3’-FAM-labeled RNA under different reaction conditions at 60 nM yDcpS. All reactions contained an additional 30 mM NaCl because of the contribution from the enzyme storage buffer. **A**) The effect of salt on the reaction. The yDcpS reaction buffer, which is 10 mM Bis-Tris pH 6.5 and 1 mM EDTA, was supplemented with the indicated concentration of salt. **B**) The effect of pH on the reaction. The concentration of each buffering agent was 10 mM and contained 1 mM EDTA. The VCE Buffer is from NEB. See Methods section for more details.

| Supplemented Salt | % decapped |
| --- | --- |
| 0 mM NaCl | >95 |
| 5 mM NaCl | >95 |
| 50 mM NaCl | >95 |
| 100 mM NaCl | 42 |
| 200 mM NaCl | 6 |
| 5 mM KCl | >95 |
| 50 mM KCl | 72 |
| 100 mM KCl | 20 |
| 200 mM KCl | <5 |

B
| Buffer, pH | % decapped |
| --- | --- |
| NaOAc pH 5.5 | 33 |
| NaOAc pH 6 | 79 |
| Bis-Tris pH 6.5 | 82 |
| K phosphate pH 7 | 16 |
| Tris-HCl pH 7.5 | 35 |
| Tris-HCl pH 8 | 13 |
| Tris-HCl pH 8.5 | <5 |
| VCE Buffer pH 8 | <5 |

### The effect of RNA 5’ secondary structure on decapping accessibility

To determine whether the substrate secondary structure has an impact on the yDcpS activity, complementary synthetic RNA oligos were annealed to the 3’-FAM-labeled 25mer Cap 0 RNA to create a blunt end, a 10-nucleotide 5’ recessed cap, or a 5-nucleotide 5’ extended cap end. The extent of decapping was compared to the single-stranded capped 25mer. All 3 double-stranded substrates were decapped, with the 5’ extended cap substrate being decapped as efficiently as the single-stranded control, while the 5’ recessed and blunt ended caps were more resistant to decapping. These results indicate that while yDcpS prefers unstructured 5’ ends, structured ends can also be decapped at higher enzyme concentration (Figure 3).

**Figure 3.**
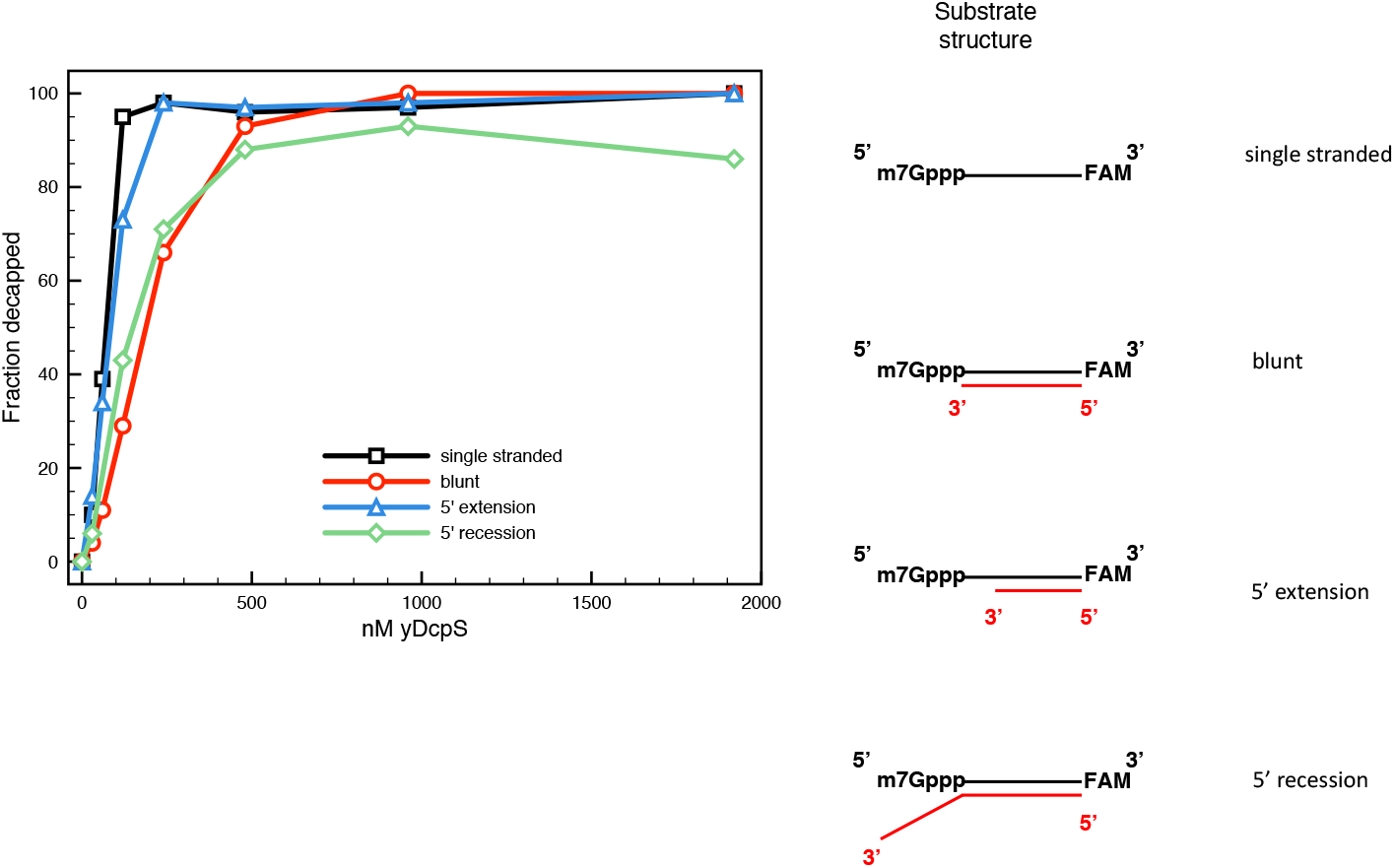
The effect of the secondary structure at the 5’ end of RNA on the decapping activity of yDcpS. Complementary synthetic RNA oligos were annealed to the 3’-FAM-labeled 25mer Cap 0 RNA to create a blunt (○), a 5’ extension (△), or a 5’ recession (⬦) as depicted on the right. The extent of decapping of each after 60 minutes at 37 °C with various concentrations of yDcpS was determined by capillary electrophoresis. The following complementary sequences were used for blunt, UUGAGCGUACUCGACGAAGUUCUAC; 5’ extension UUGAGCGUACUCGACGAAGU; and 5’ recession, UUGAGCGUACUCGACGAAGUUCUACAAUGACCAUC.

### Cap specificity of yDcpS

Liu *et al.*^4^ suggested that the cap analog GpppG was not likely a substrate for DcpS because the GpppG dinucleotide at 10 μM was not an effective inhibitor of m^7^GpppG decapping. Wypijewska *et al.*^10^ and Cohen *et al*.^11^ demonstrated that DcpS from *H. sapiens*, *C. elegans*, and *A. suum* differ from one another as they decap various cap dinucleotide analogs at dissimilar rates. Yeast mRNA caps are only methylated at the *N*-7 position and not at the 2’-*O* position^19,20^. In contrast, the majority of the capped structures from HeLa cells are Cap 1 (m^7^GpppNm) and Cap 2 (m^7^GpppNmNm), whereas Cap 0 (GpppN) has not been identified^21,22^. Additionally, a number of still uncharacterized cap structures have been reported to occur on short RNAs ^23^. Hence, we sought to determine the susceptibility of 30 distinct 25mer RNAs featuring different cap structure permutations to decapping with yDcpS (Figure 4a).

**Figure 4.**
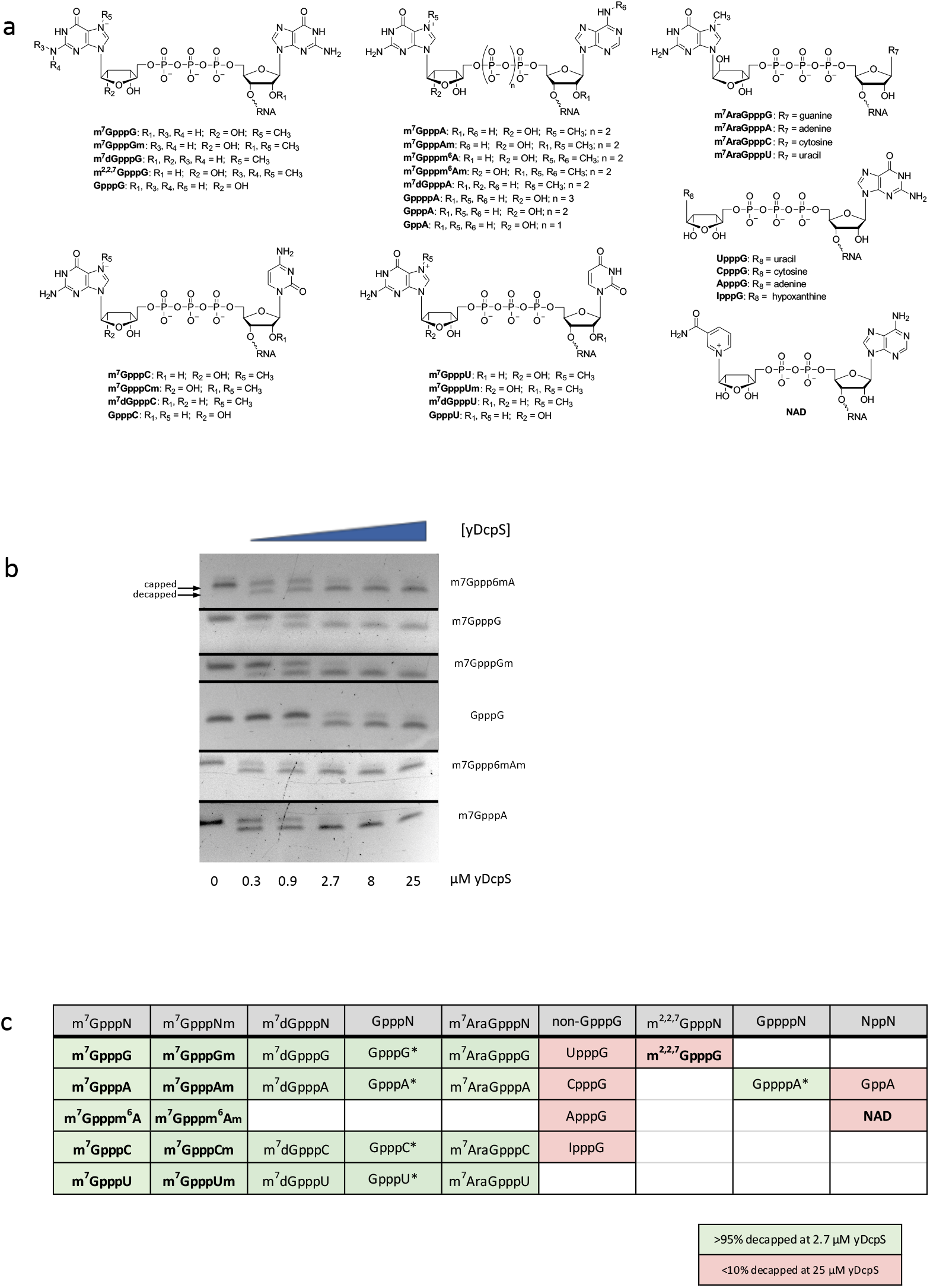
Relative decapping efficiency of yDcpS with 30 different cap structures on 25mer RNAs. (**a**) chemical representation of cap structures. (**b**) Composite of gel images of 6 decapping reactions of 25mer RNA substrates (m^7^Gppp^6^mA, m^7^GpppG, m^7^GpppGm, GpppG, m^7^Gppp^6^mAm, m^7^GpppA) incubated with increasing concentrations of yDcpS for 60 minutes at 37 ̊C. The reactions were electrophoresed on a 15% TBE-Urea gel and stained with SYBR Gold. The full-length gel images are shown in the Supplemental Information. (**c**) Substrates decapped (greater than 95% at 2.7 μM yDcpS) are shaded in green and substrates resistant to decapping (less than 10% at 25 μM yDcpS) are shaded in pink. Bolded characters indicate canonical cap structures (known or anticipated). Asterisks indicate 5-10 fold reduced decapping relative to m7GpppG-25mer at 0.9 μM yDcpS.

Various capped 25mer RNA substrates (100 nM) were incubated for 60 minutes at 37 °C with increasing concentrations of yDcpS varying by a factor of three. The reactions were then analyzed by PAGE and the relative extent of decapping determined after staining and imaging the gels. A typical set of gel images was compiled in Figure 4b. It can be estimated from the gel electrophoresis results that, m^7^GpppA, m^7^Gpppm^6^A, m^7^Gpppm^6^Am, m^7^GpppG, and m^7^GpppGm caps were removed at a similar yDcpS concentration of 0.9 μM, while GpppG required about a 3-fold higher concentration. Interestingly, while yDcpS can decap deoxyriboguanosine and arabinoguanosine caps, no activity was detected towards 2,2,7-trimethylguanosine, adenosine, cytidine, or uridine capped RNAs (Figure 4c). As for the length of the internucleotidic phosphate bridge, yDcpS was able to decap tri- or tetraphosphate, but no diphosphate caps. Not surprisingly, yDcpS had no activity on a nicotinamide adenine dinucleotide (NAD) capped RNA.

### yDcpS decapping followed by VCE recapping

A characteristic of yDcpS catalysis is the removal of the m^7^G from the cap in the form of m^7^GMP leaving a 5’-diphosphate on the 5’ terminus of the RNA^4^. 5’-diphosphate RNA is a known substrate for *in vitro* capping using the *Vaccinia* virus Capping Enzyme (VCE)^9,24,25^. We took advantage of these two reactions to tag a m^7^G-capped 25mer RNA with an affinity group, by first decapping the RNA with yDcpS, and then recapping it by treatment with VCE and DTB-GTP^9^. These reactions were performed in the presence of excess total *E. coli* RNA in order to simulate a complex mixture of RNA as would be found in total eukaryotic RNA. As shown by gel electrophoresis for a Cap 1 RNA (Figure 5) and by mass spectroscopy for Cap 0 (Supplementary Fig. S1-S3), the decapping/recapping process approached completion, suggesting that this strategy could be extended to capture and enrich native capped RNA transcripts.

**Figure 5.**
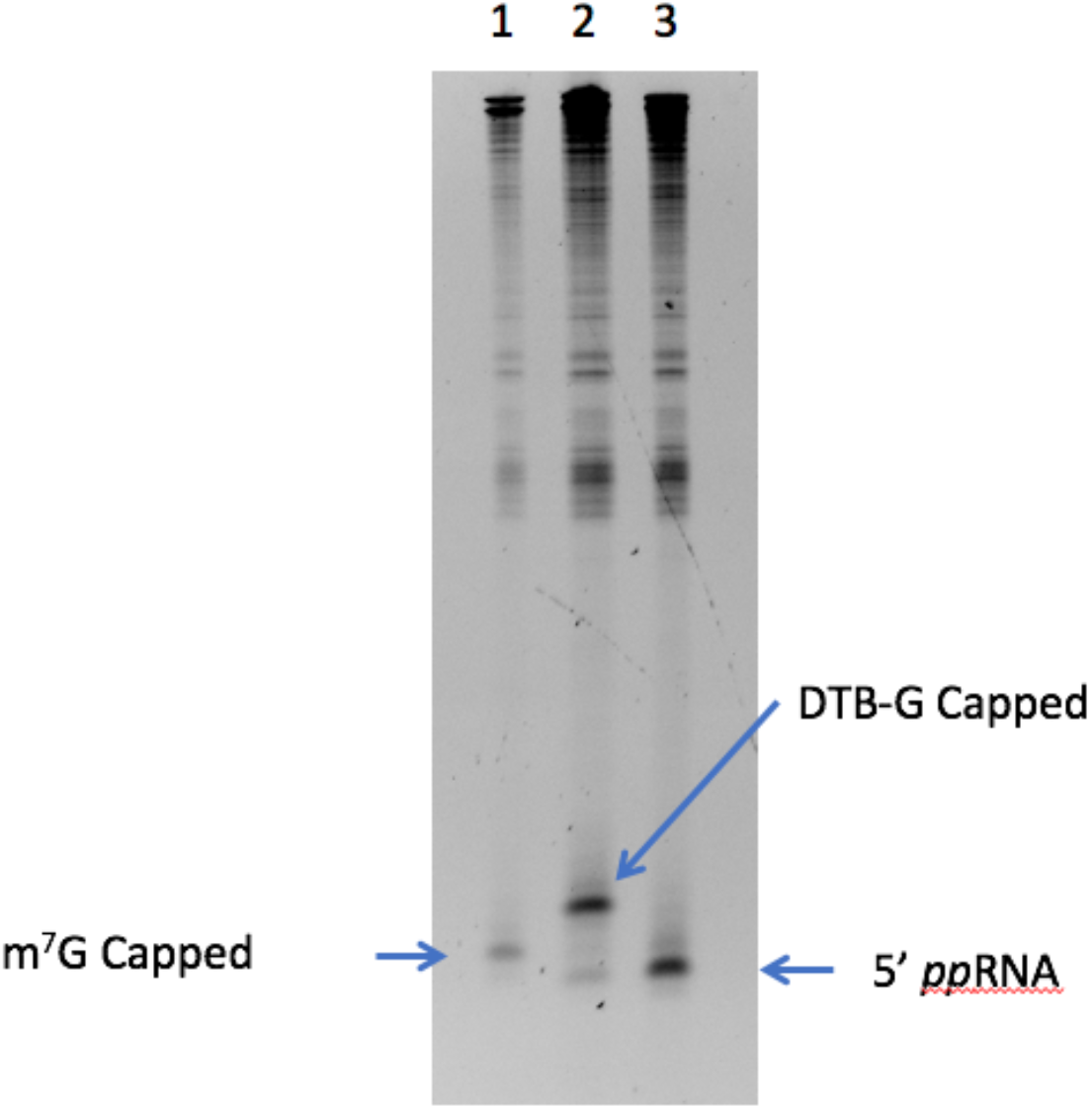
yDcpS decapping of a 25mer Cap1 RNA followed by VCE recapping. A m^7^G-capped 25mer RNA (0.22 μg) was mixed with 1 μg of total *E. coli* RNA and incubated with 0.1 nmol of yDcpS for 2 h at 37 °C. An aliquot was removed prior to the addition of yDcpS to provide a reference band for m^7^G-capped 25mer (Lane 1). The yDcpS reaction was terminated by addition of Proteinase K and purified using Ampure XP beads. After elution, the RNA was incubated in 1X VCE buffer containing 0.1 mM SAM and 0.5 mM DTB-GTP in the presence (Lane 2) or absence (Lane 3) of VCE for 1 h at 37 °C. All samples were electrophoresed on a 15% TBE-Urea gel and stained with SYBR Gold.

### yDcpS decaps long capped RNA transcripts

After demonstrating the efficient decapping and recapping of m^7^G-capped 25mer RNAs, we tested whether yDcpS could decap long RNA substrates. We generated a mixture of RNA transcripts with lengths of 90 to 1400 nucleotides by *in vitro* transcription from a plasmid harboring a T7 promoter upstream of the FLuc gene, which had been cleaved with various restriction endonucleases to generate transcription templates of different lengths (see Methods). The produced RNA transcripts were capped with GTP by VCE to yield m^7^G-capped RNAs. The mixture of capped transcripts was then decapped by incubation with yDcpS. As a control, an aliquot of the m^7^G-capped transcripts was incubated in the absence of yDcpS. Both sets of transcripts were subjected to capping with DTB-GTP by VCE. The recapped products were exposed to streptavidin beads, and after washing steps to remove any unbound material, recovered by elution with biotin. As shown in Figure 6, no RNA was recovered from the fraction that was not treated with yDcpS (Lane 4), while the full-size range of capped transcripts treated with yDcpS were recovered in the biotin-eluted fraction. This result demonstrates that yDcpS decapping followed by VCE recapping and streptavidin-based recovery of transcripts is an efficient process (approximately 50% yield) and shows minimal length discrimination. This workflow can thus be applied to eukaryotic mRNA Transcription Start Site (TSS) analysis by appending it to the workflow of Cappable-seq^9^.

**Figure 6.**
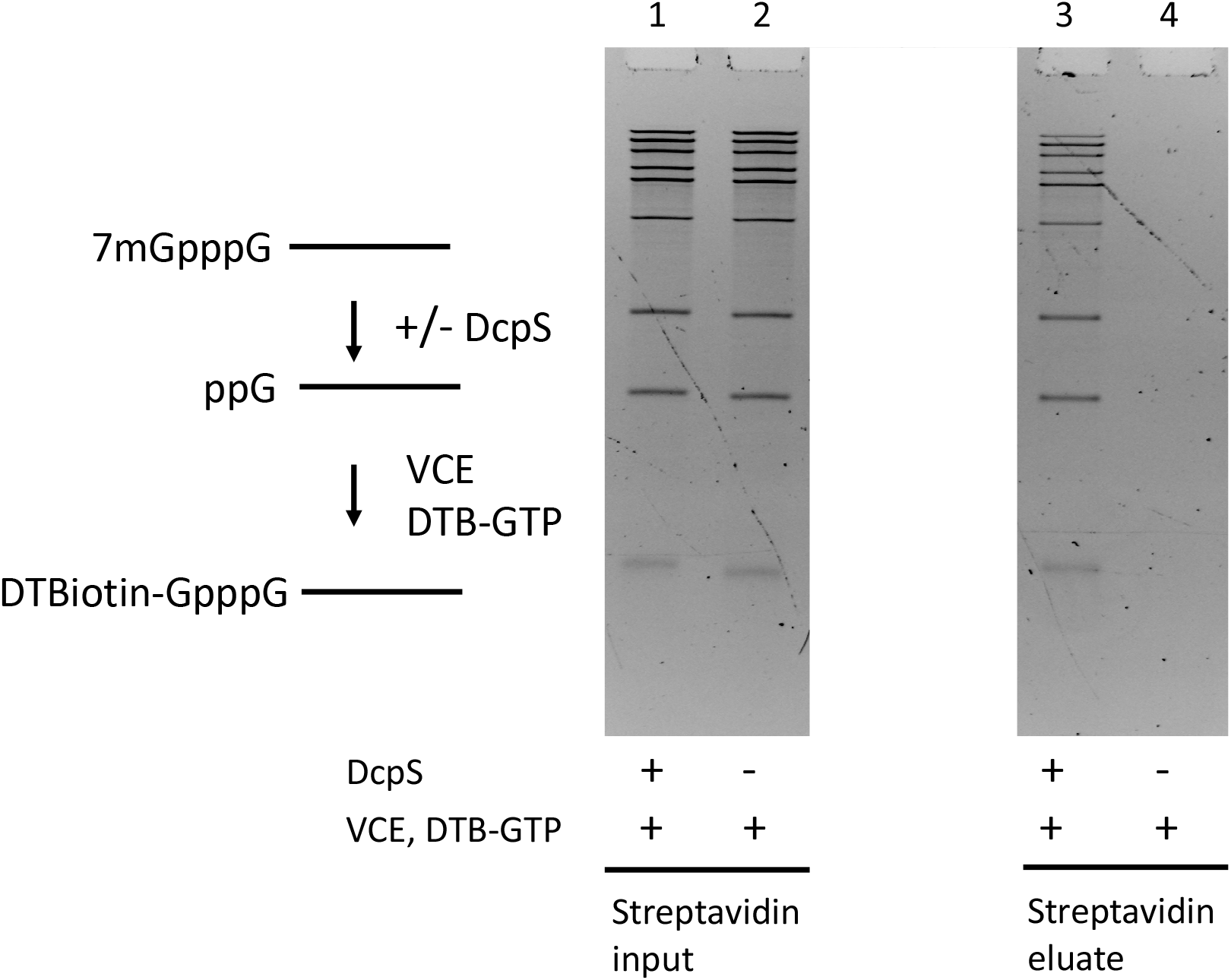
Efficient recovery of long capped RNA transcripts by a decapping/recapping procedure. A mixture of m^7^G-capped RNAs (0.090 -1.4 kb) was divided into two samples. One sample was incubated with yDcpS and one without yDcpS, and both were subsequently treated with VCE and DTB-GTP. An equal fraction of each was exposed to streptavidin beads, which were washed and eluted. Lanes 1 (+ yDcpS) and 2 (− yDcpS) show the samples after the VCE reaction. Lanes 3 (+ yDcpS) and 4 (− yDcpS) show the eluates from streptavidin beads. All lanes represent an equal fraction of the original mixture. Both gel panels are from the same imaged gel.

## Discussion

In characterizing yDcpS, we have shown that the methylation status of various canonical guanosine caps (GpppN, m^7^GpppN, m^7^GpppNm, m^7^Gpppm^6^A, or m^7^Gpppm^6^Am) does not significantly affect their susceptibility to decapping by this enzyme. We also examined non-canonical m^7^G caps where the ribose moiety was substituted with arabinose or deoxyribose. Although these caps have not been shown to exist in nature, they are also decapped by yDcpS.

Under the conditions described, yDcpS decaps the majority of the guanosine-based caps, regardless of the nucleotide at position +1. The enzyme prefers m^7^G over nonmethylated G caps, and does not decap adenosine, cytosine, uridine, or inosine caps. yDcpS decaps 5’-5’ triphosphate and tetraphosphate linkages, but not diphosphate linkages. Interestingly, Cohen *et al.*^11^ reported that nematode DcpS decapped a 2,2,7-trimethylated G cap (commonly found in certain small nuclear and nucleolar RNAs), while the human DcpS did not. We found that yDcpS, resembling hDcpS, did not decap 2,2,7-trimethylated guanosine.

The *in vitro* characterization of yDcpS presented here provides evidence that yDcpS should be a useful reagent for generating recappable 5’ ends from capped RNAs. We demonstrate that yDcpS decaps m^7^G-capped RNAs from as short as 25 to at least 1400 nucleotides, leaving a 5’-diphosphate end. The process of decapping and recapping with an affinity-tagged nucleotide would be advantageous for identifying and enriching 5’ capped termini of mRNAs in eukaryotes. This would expand the utility of the prokaryotic Cappable-seq method^9^ to include eukaryotic mRNAs. Cappable-seq has been used to determine the transcription start sites of primary transcripts of prokaryotes at single base resolution via the attachment of a biotinylated GTP analogue to the 5’-triphosphate end of primary RNA transcripts by action of the *Vaccinia* capping enzyme (VCE). The extension of this method to canonical eukaryotic mRNAs requires the removal of the m^7^G moiety to generate a recappable end (such as a tri- or diphosphate 5’ end), which we now show is attainable with the use of yDcpS. Therefore, we anticipate being able to enrich eukaryotic mRNA by combining yDcpS-mediated decapping followed by recapping with affinity tagged GTP and generating libraries for high-throughput sequencing. Progress towards this goal is underway in our laboratory.

## Methods

### Preparation of RNAs

The m^7^GpppA dinucleotide cap analog m^7^G(5’)ppp(5’)A was obtained from New England Biolabs, Ipswich, MA (NEB). The 5’-[m^7^Gppp]GUAGAACUUCGUCGAGUACGCUCAA[FAM]-3 was purchased from Bio-Synthesis, Inc. The complementary oligos used for secondary structure experiments for blunt end (UUGAGCGUACUCGACGAAGUUCUAC), 5’ cap extension (UUGAGCGUACUCGACGAAGU), and 5’ cap recession (UUGAGCGUACUCGACGAAGUUCUACAAUGACCAUC) were purchased from Integrated DNA Technologies (IDT). The 5’-triphosphate 25mer RNAs (5’-*ppp*-NUAGAACUUCGUCGAGUACGCUCAA-3’, wherein N is G, C, A, or U) utilized for the generation of m^7^GpppN, m^7^GpppNm, m^7^dGpppN, GpppN, and m^7^AraGpppN were individually synthesized as previously described^26^. The 5’-monophosphate 25mer RNAs (5’-*p*-NUAGAACUUCGUCGAGUACGCUCAA-3’, wherein N is G, A, or U) utilized for the generation of m^7^Gpppm^6^A, m^7^Gpppm^6^Am, GppppA, GppA, and NpppG (wherein N is U, C, A, or I) were synthesized by standard phosphoramidite chemistry on an ABI 394 DNA/RNA synthesizer. For the synthesis of 5’-*p*-m^6^AUAGAACUUCGUCGAGUACGCUCAA-3’, a *N*6-Methyl-A-CE phosphoramidite from Glen Research was utilized. Detailed protocols for the enzymatic or chemical generation of the 5’-capped 25mer RNAs utilized in this study are described in the Supplementary Information section.

### Preparation of yDcpS

The gene for the *Saccharomyces cerevisiae* yDcpS (YLR270W) was codon optimized for *E. coli* expression and designed with an amino terminal His tag to be expressed from a T7 promoter in plasmid pET28a (Supplementary Fig. S4). The predicted molecular weight of the fusion is 42,945 daltons. The construct was synthesized by Genscript. The protein was expressed in *E. coli* ER3600 and purified to near homogeneity (greater than 95%) by chromatography over HisTrap, SP, and Q resins. The protein’s size as determined by SDS-PAGE is consistent with its predicted molecular weight. The final preparation of yDcpS was 12 μg of protein/μL, in a storage buffer of 20 mM Tris-HCl pH 7.5, 200 mM NaCl, 0.1 mM EDTA, and 50% glycerol. The protein was stored at −20 °C. The DcpS preparation was determined to be free of detectable RNAse contamination by assay with a 300mer RNA transcript (~100 ng) incubated with 12 μg of yDcpS in a 20 μL reaction in the decapping buffer 10 mM Bis-Tris pH 6.5, 1 mM EDTA, and in 20 mM Tris-Acetate pH 7.9 buffer containing 50 mM potassium acetate, 10 mM magnesium acetate, and 1 mM DTT for 4 hours. No degradation of the 300mer band by PAGE analysis was observed.

### Human DcpS

Human DcpS (hDcpS) was purchased from Enzymax LLC, Lexington, KY. The protein concentration of DcpS was determined by OD_280_ using the molar extinction coefficient of 30495 M^−1^ cm^−1^.

### Decapping reactions

Unless otherwise noted, the decapping was carried out at 37 °C in the decapping buffer 10 mM Bis-Tris pH 6.5, 1 mM EDTA. With the exception of the reactions containing FAM-labeled RNAs (see below), all reactions using capped 25mer RNA substrates were terminated by the addition of 1 volume of 2X RNA loading dye (NEB). The reactions using the m^7^GpppA dinucleotide cap analog were terminated with the addition of phenol/chloroform.

### *In vitro* decapping of 5’-capped synthetic 3’-FAM-labeled RNA

*In vitro* decapping reactions were carried out in a 20 μL reaction containing 10 mM Bis-Tris pH 6.5, 1 mM EDTA, 500 nM substrate RNA (5’-[m^7^Gppp]GUAGAACUUCGUCGAGUACGCUCAA[FAM]-3, Bio-Synthesis, Inc.), and 60 nM yDcpS at 37 °C for 60 minutes, unless otherwise indicated (Table 1). Reactions were stopped by heat inactivation at 70 °C for 15 minutes. Reactions were diluted in nuclease-free water to reach a final substrate concentration of 5 nM before capillary electrophoresis on either an Applied Biosystems 3130xl Genetic Analyzer (16 capillary array) or an Applied Biosystems 3730xl Genetic Analyzer (96 capillary array) using GeneScan 120 LIZ dye Size Standard (Applied Biosystems). Reaction products were analyzed using PeakScanner software (Thermo Fisher Scientific) and an in-house software suite. The electrophoretic mobility of the Cap 0 substrate and the 5’-diphosphate product was calibrated against RNA standards generated enzymatically from a synthetic 5’-triphosphate 25mer. The 5’-diphosphate standard was generated by treating the chemically synthesized 5’-triphosphate/3’-FAM-labeled 25mer with the *S. cerevisiae* Cet1^27^. The 5’-monophosphate standard was generated by treating the 5’-triphosphate/3’-FAM-labeled 25mer with Apyrase (NEB). The 5’-hydroxyl standard was generated by treating the 5’-triphosphate/3’-FAM-labeled 25mer with calf intestinal phosphatase (NEB). The cap 0 standard was generated by treating the 5’-triphosphate/3’-FAM-labeled 25mer with *Vaccinia* virus capping enzyme (VCE, NEB) in the presence of GTP and *S*-adenosylmethionine (SAM). The RNA standards were purified using RNA Clean and Concentrator kit (Zymo Research). The identity of the standards was verified by intact mass spectrometry. The 25mer Cap 1 RNA was generated by treating the chemically synthesized 3’-FAM-labeled Cap 0 25mer RNA with mRNA Cap 2’-*O*-Methyltransferase (NEB) in the presence of SAM. The Cap 1 25mer RNA was purified using Monarch RNA Cleanup kit (50 μg capacity, NEB). Complete methylation was verified by digestion of RNA with the Nucleoside Digestion Mix (NEB, M0649) and LC-MS analysis.

### *In vitro* cap specificity assay of yDcpS with 30 different cap structures

The reaction conditions for decapping consisted of 100 nM 5’-capped 25mer RNA and 0, 0.3, 0.9, 2.7, 8 or 25 μM DcpS in a 10 μL reaction volume in 10 mM Bis-Tris pH 6.5 and 1 mM EDTA. The reactions were incubated for 60 minutes at 37 °C and terminated by the addition 2 μL of Proteinase K (NEB) which had been supplemented with 0.1 reaction volume of 1 M Tris-HCl pH 8.0, and further incubated for 5 minutes at room temperature. To each 12 μL reaction was added 12 μL of RNA loading dye (2X) (NEB). This mixture was heat denatured at 80 °C for 3 minutes, and an aliquot of 5 μL was loaded onto a 15% polyacrylamide TBE-Urea gel (Invitrogen). The gel was run at 180 volts for 75 minutes. The RNA was visualized with SYBR Gold stain.

### Decapping 25mer with yDcpS and recapping with 3’-Desthiobiotin-GTP

A 100 μL reaction in 10 mM MES pH 6.5 and 1 mM EDTA containing 0.22 μg of 25mer Cap1 RNA and 0.1 nmol of yDcpS, and 1 μg of total *E. coli* RNA, was incubated for 2 hours at 37 °C. Before addition of the yDcpS, a 2 μL sample of the reaction was removed. The yDcpS reaction was terminated by addition of 4 μL of 1M Tris HCl pH 8.0 and 2 μL of Proteinase K. The reaction was further incubated for 30 minutes at 37 °C and then heated to 94 °C for 3 minutes. The RNA was purified by binding to Ampure XP beads (Beckman Coulter) by addition of 2 volumes of beads, and to that final volume additional 1.5 volumes of ethanol. After washing the beads with 80% ethanol, the RNA was eluted in 30 μL of 1 mM Tris pH 7.5 and 0.1 mM EDTA. A 20 μL reaction containing 1X VCE buffer (NEB), 10 μL of the eluted RNA, 0.1 mM SAM, 0.5 mM DTB-GTP (NEB) was split into two 9.5 μL reactions: in one tube was added 1 μL of VCE and in the other tube was added 1 μL of water. Samples were electrophoresed on a 15% TBE-Urea gel (Invitrogen) and stained with SYBR Gold. A similar experiment for decapping and recapping was carried out with a Cap 0 25mer RNA. The results of this experiment were confirmed by mass spectrometry (Supplementary Fig. S1-S3).

### Preparation of long (0.09 - 1.4 kb) m^7^G-capped transcripts

m^7^G-capped RNA substrates 90 to 1400 nucleotides long were generated by T7 RNA polymerase (NEB) transcription of different DNA templates generated from an FLuc plasmid, each digested with one of the following restriction endonucleases: NlaIV, HpyCH4V, Hpy166II, PsiI, AflIII, EcorI, BspQI, BcoDI, or EcoRV in separate reactions. The resulting RNA transcripts comprised 90, 171, 240, 419, 587, 681, 896, 1117, and 1432 nucleotides, respectively. The individual transcripts were combined in approximate equal amounts. The transcript mixture was subsequently capped by standard VCE capping reaction with GTP and SAM.

### Data acquisition and settings

RNA was loaded onto 6% or 15% PAGE Urea gels and electrophoresed for 75-85 minutes at 180 V. The gels were stained with SYBR Gold and imaged with an AlphaImager HP. The images were imported into Photoshop, inverted and levels were auto-adjusted.

## Acknowledgements

We thank New England Biolabs for supporting this work.

## Author Contributions

I.S. conceived and designed the study. I.S. and I.R.C.J. wrote the manuscript with input from all the authors. M.W. and J.B. synthesized the majority of variously capped RNAs. E.M., G.T., J.W. and I.S. performed biochemical assays. K.M. and I.S. purified the proteins. S-H.C. developed the CE assays and N.D. performed LC-MS analysis. All authors approved the manuscript.

