## Supplementary file for "*In vitro* characterization of the yeast DcpS, a scavenger mRNA decapping enzyme"

### Preparation of Cap 0 and Cap1 25mer RNAs by T7-mediated transcription

A T7 transcription reaction using a synthetic double-stranded DNA template was carried out using the HiScribe T7 High Yield RNA Synthesis Kit (NEB) to yield the 25mer transcript 5'-*ppp*GGGAGTCTTCGTCGAGTACGCTCAA. The transcript was purified by a Clean and Concentrator spin column (Zymo Research). The RNA was capped using GTP and SAM with the *Vaccinia* Capping System (NEB). The RNA was purified with Clean and Concentrator (Zymo Research) to produce the Cap 0 25mer RNA. The Cap 0 RNA was converted to Cap 1 with mRNA Cap 2'-O-Methyltransferase (NEB).

### Preparation of synthetic 5'-capped 25mer RNAs by enzymatic capping

For the generation of $m^7$GpppN, $m^7$GpppNm, $m^7$dGpppN, GpppN, and $m^7$AraGpppN, 5'-triphosphate RNAs (5'-*ppp*-NUAGAACUUCGUCGAGUACGCUCAA-3', wherein N is G, C, A, or U) were prepared according to published methods[1]. Enzymatic capping of 5'-triphosphate RNAs (5 nmol) was performed at a 500 µL reaction volume using the Vaccinia Capping System (NEB) supplemented with yeast inorganic pyrophosphate (NEB) as follows: 10 µM 5'-triphosphate RNA, 50 mM Tris-HCl pH 8, 5 mM KCl, 1 mM $MgCl_2$, 1 mM DTT, 30 µM GTP, 200 µM SAM , 50 units of pyrophosphatase, and 500 units of VCE. To obtain non-methylated GpppN caps, SAM was omitted from the reaction. For $m^7$dGpppN and $m^7$AraGpppN caps, the corresponding dGTP and AraGTP used at 100 µM. For Cap 1 structures ($m^7$GpppNm), 2500 units of mRNA Cap 2'-*O*-Methyltransferase (NEB) were included in the reaction. Prior to the capping reactions, the RNA dissolved in water was heat-denatured at 65 °C for 5 minutes and placed on ice for an additional 5 minutes. The remaining reaction components were added, and the reaction was allowed to proceed at 37 °C overnight. The enzyme and small molecules were removed from the reaction using phenol/chloroform extraction with Phase Lock Gel tubes (5Prime) following the manufacturer's protocol. The aqueous phase containing the capped oligonucleotides was then transferred to in a fresh tube and concentrated on a SpeedVac (Savant).

The crude material was purified using 20% polyacrylamide gel electrophoresis (PAGE) in TBE 1X buffer. Immediately prior to purification, the samples were resuspended in 50 µL of 7 M urea and heat-denatured at 70 °C for 5 minutes. The gel was run at 800 V constant voltage for up to 6 h. The capped oligonucleotides were revealed by UV shadowing against a silica plate with fluorescent indicator. The band of interest was cut out and crushed in 2 mL centrifuge tubes with disposable plastic pestles, resuspended and vortexed in 500 µL of 500 mM Tris pH 7.5. The gel particles were heated at 60 °C for 15 minutes, immediately frozen at -80 °C for 30 minutes, and then left overnight at room temperature. The gel solution was centrifuged at 15,000 rpm for 30 minutes, and the supernatant was removed and filtered with Ultrafree-MC 0.45 µm tubes (Millipore). An additional 500 µL of 500 mM Tris pH 7.5 was added to the particles, vortexed, and centrifuged again. The recovered supernatant fractions were pooled and precipitated with 100 µL of 3 M sodium acetate in 2 mL of 100% absolute ethanol. The mixture was vortexed and frozen for at least 30 minutes at -80 °C. The tube was centrifuged for 30 minutes at 15,000 rpm, and the solution was discarded. The pellet was washed with 500 µL of 70% ethanol and centrifuged again. The sample was vacuum-dried using a SpeedVac Concentrator. The pellet of purified oligonucleotide was resuspended in 50 µL of water and stored at -20 °C. The final capped oligonucleotide concentration was estimated with a NanoDrop spectrophotometer. The intact mass of all capped oligonucleotides was confirmed by mass spectrometry (Oligo HTCS, Novatia LLC).

### Preparation of synthetic 5'-capped 25mer RNAs by chemical capping

For the generation of $m^7Gpppm^6A$, $m^7Gpppm^6Am$, GppppA, GppA, and NpppG (where N is U, C, A, or I), 5'-monophosphate 25mer RNAs (5'-*p*-NUAGAACUUCGUCGAGUACGCUCAA-3', wherein N is G, A, or $m^6A$) were made by standard solid-phase synthesis. Chemical capping of 5'-monophosphate RNAs (5 nmol) was carried out at a 250 µL reaction scale according to method adapted from Sawai *et al.*[2]. Briefly, 50 µL of 100 µM 5'-monophosphate RNA was combined with 50 µL of 1 M Bis-Tris buffer pH 6.0, 5 µL of 1 M $MnCl_2$ , and 50 µL of DMF . To this solution was added 95 µL of a 100 mM solution of corresponding imidazolide-NMP,

-NDP, or -NTP, and the reaction was incubated at 50 °C for 5 hours, 37 °C for 5 hours, or room temperature for 4 hours for GppA, NpppG, or GppppA 25mer RNAs, respectively. After this time, the unreacted imidazolide was removed from the reaction along with salts and organic solvent using Sep-Pak C18 Cartridges (Waters WAT051910) as follows: the capping reaction was diluted to 5 mL in 0.1 M triethylammonium bicarbonate (TEAB), pushed through the Sep-Pak column and washed with an additional 15 mL of 0.1 M TEAB. The capped oligonucleotide was eluted from the column using 1:1 TEAB:Acetonitrile (2 mL). The capped oligonucleotide was concentrated on a SpeedVac. To generate $m^7Gpppm^6A$ or $m^7Gpppm^6Am$ from the chemically capped $Gpppm^6A$ 25mer RNA, the crude material was subjected to an enzymatic treatment with VCE and SAM as described above. To generate $m^7Gpppm^6Am$ from $m^7Gpppm^6A$, the crude material was further treated with mRNA Cap 2'-*O*-Methyltransferase. All 5'-capped 25mer RNAs were purified by polyacrylamide gel electrophoresis described above. Their intact mass was confirmed for by mass spectrometry (Oligo HTCS, Novatia LLC).

**Preparation of 2,2,7-trimethylated guanosine cap 25mer RNA**

The $m^7GpppG$ 25mer RNA was converted to the trimethylated form $m^{2,2,7}GpppG$ by incubation with the human trimethylguanosine synthase 1 (hTGS)[3]. A 100 µL reaction consisting of 50 pmol of $m^7GpppG$-25mer, 150 µM SAM, 50 mM Tris-HCl pH 7.5, 1 mM EDTA, and 200 ng of a hTGS preparation for 2 hours at 37 ˚C. The product was purified by spin column with a Monarch RNA Cleanup Kit (10 µg capacity, NEB). The resulting trimethylguanosine cap 25mer RNA product was analyzed by intact mass spectroscopy (Oligo HTCS, Novatia LLC) (Supplementary Figure 5).

The hTGS enzyme preparation was obtained as follows: a plasmid harboring a truncated hTGS gene (618-853) as a carboxy terminal fusion of the maltose binding protein (MBP) gene was obtained from Genscript USA. The plasmid was induced with IPTG and incubated at 18 ˚C overnight. The fusion protein was purified by amylose chromatography followed by heparin sepharose chromatography.

#### Preparation of 5'-NAD capped 25mer RNA

The 5'-NAD capped 25mer was generated by cotranscriptional incorporation of nicotinamide adenine dinucleotide (NAD) at the initiating nucleotide position. A 10 µM mixture of the template (TTGAGCGTACTCGACGAAGTTCTCTAATAGTGAGTCGTATTAGCTTCTGTAC) and non-template strand (GTACAGAAGCTAATACGACTCACTATTAGAGAACTTCGTCGAGTACGCTCAA) DNA oligos (IDT) was heated to 90 °C for 30 s and allowed to cool slowly to room temperature. Transcription was carried out using HiScribe T7 High Yield RNA Synthesis Kit (NEB) supplemented with NAD, $MgCl_2$, and DMSO as follows: 1x T7 Reaction Buffer, 4 mM NAD, 2 mM each of the four NTPs, 0.2 µM template, 5% DMSO, 5 mM $MgCl_2$, and 2 µL of T7 RNA Polymerase Mix at 37 °C for 2 h. The transcription reactions were then treated with DNase I (NEB) at 37 ˚C for 1 h. The NAD-capped RNA was purified using 20% urea PAGE. Transcript sequence:NicppAGAGAACUUCGUCGAGUACGCUCAA; Nic = nicotinamide ribose.

### Supplementary Figures

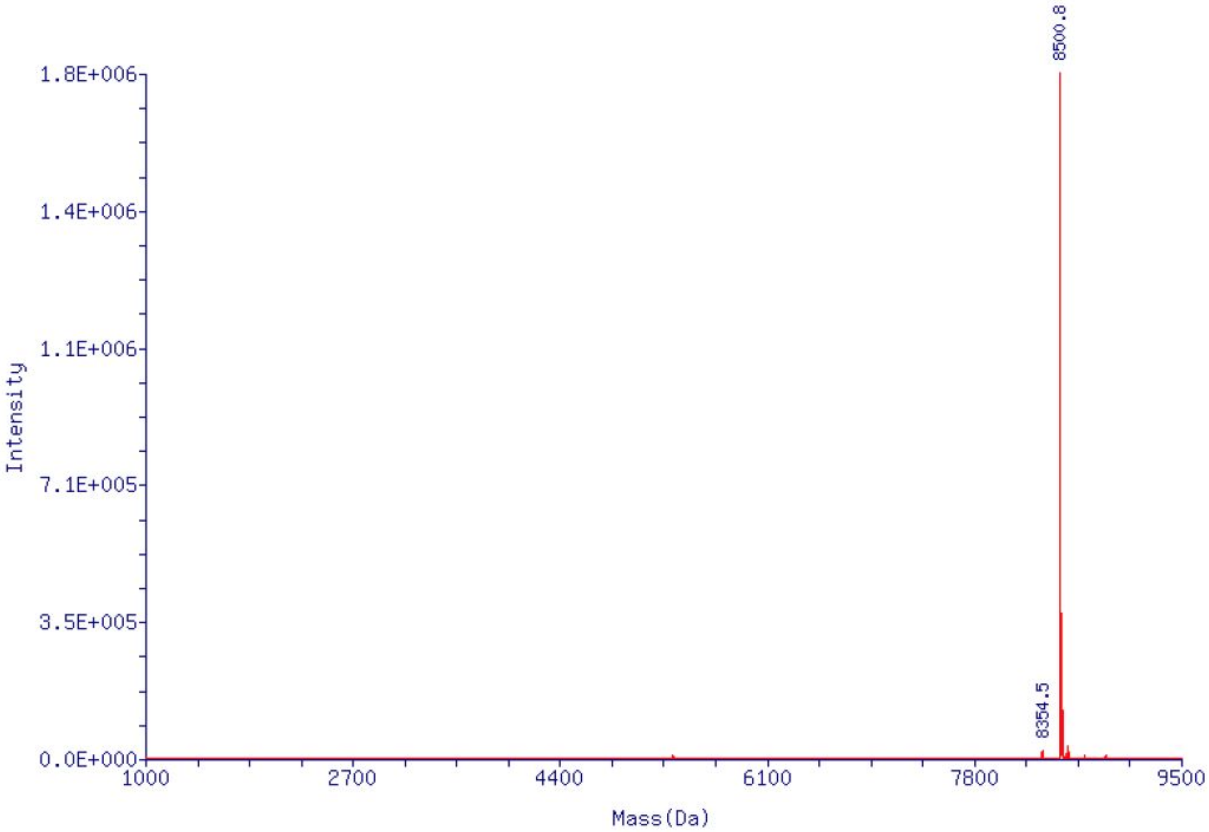


**Supplementary Figure 1.** Deconvoluted LTQ-XL mass spectrum of 5'-capped m$^7$GpppG 25mer RNA. Target Mass (Da) calculated for m$^7$G*ppp*GUAGAACUUCGUCGAGUACGCUCAA ($C_{249}H_{311}N_{100}O_{185}P_{27}$): 8501.1. Observed Mass (Da): 8500.8. Mass Error: -0.3 Da (-0.004%). Estimate Purity. 97.3%.

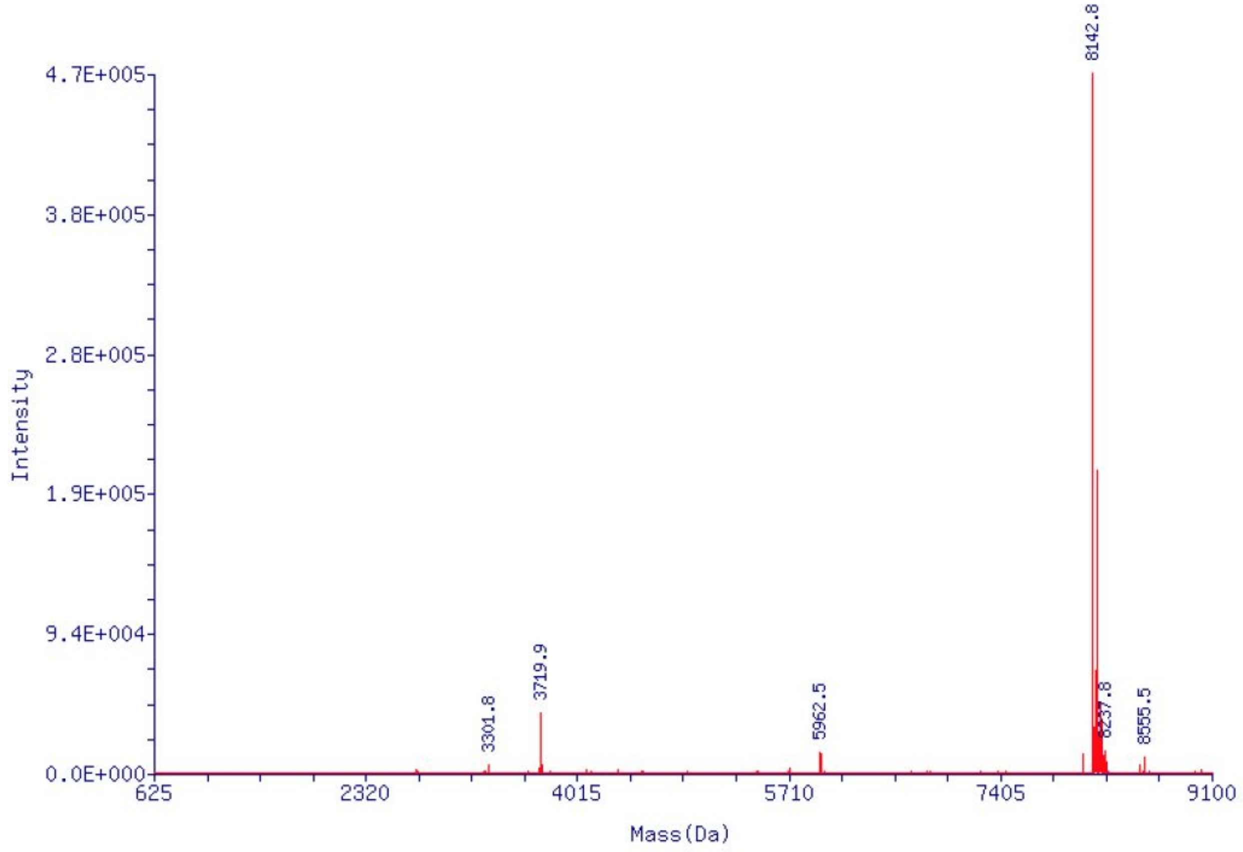


**Supplementary Figure 2.** Deconvoluted LTQ-XL mass spectrum of 5'-ppG 25mer RNA obtained from the decapping reaction of $m^7$GpppG 25mer RNA with yDcpS. Target Mass (Da) calculated for *pp*GUAGAACUUCGUCGAGUACGCUCAA ($C_{238}H_{297}N_{95}O_{178}P_{26}$): 8141.8. Observed Mass (Da): 8142.8. Mass Error: 1.0 Da (0.012%). Estimate Purity. 83.9%.

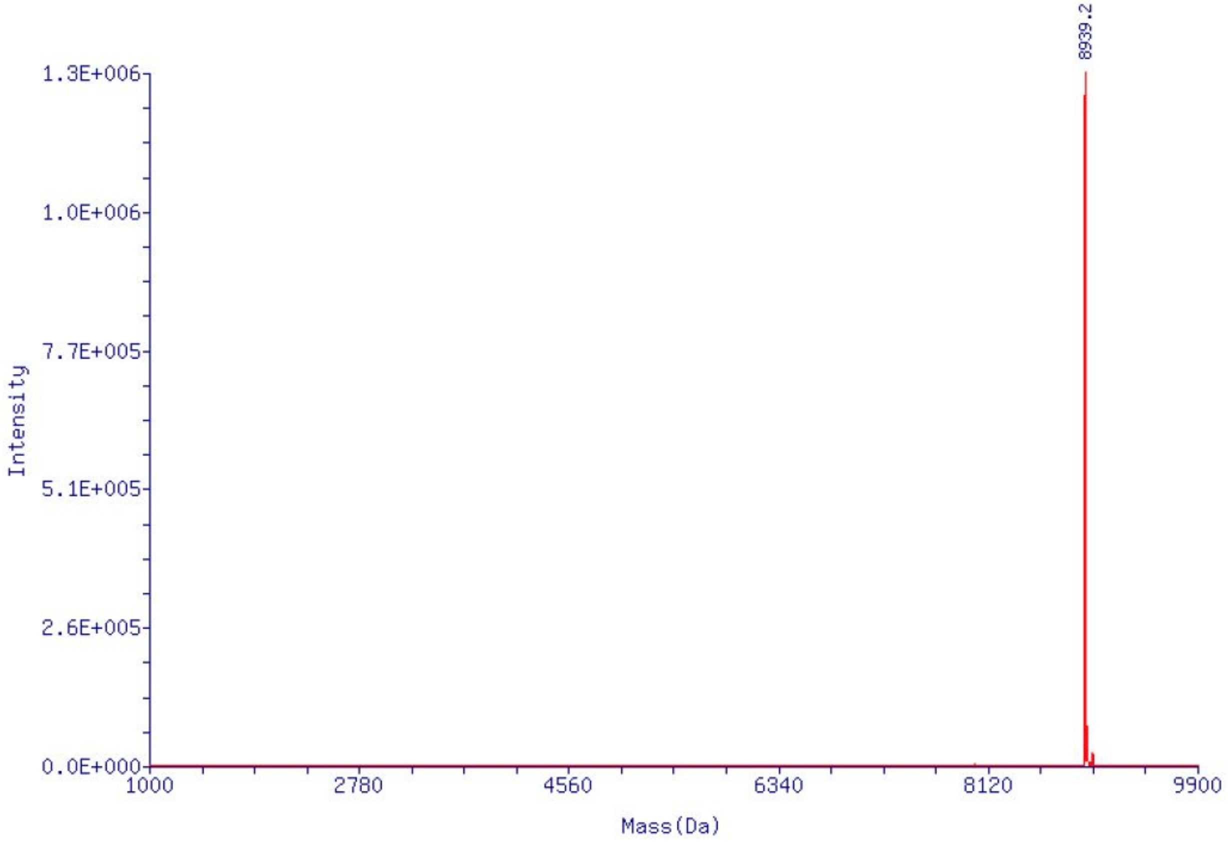


**Supplementary Figure 3.** Deconvoluted LTQ-XL mass spectrum of DBT-GpppG 25mer RNA after recapping 5'-ppG 25mer RNA (product of yDcpS treatment of Supplementary Figure 2) with VCE in the presence of 3'-Desthiobiotin-GTP. Target Mass (Da) calculated for DTB-G*ppp*GUAGAACUUCGUCGAGUACGCUCAA ($C_{269}H_{345}N_{106}O_{190}P_{27}$): 8939.6. Observed Mass (Da): 8939.2. Mass Error: -0.4 Da (-0.004%). Estimate Purity. 98.3%.

```
1         10        20        30        40        50
|         |         |         |         |         |
MGSSHHHHHHSSGLVPRGSHMSQLPTDFASLIKRFQFVSVLDSNPQTKVM
SLLGTIDNKDAIITAEKTHFLFDETVRRPSQDGRSIPVLYNCENEYSCIN
GIQELKEITSNDIYYWGLSVIKQDMESNPTAKLNLIWPATPIHIKKYEQQ
NFHLVRETPEMYKRIVQPYIEEMCNNGRLKWVNNILYEGAESERVVYKDF
SEENKDDGFLILPDMKWDGMNLDSLYLVAIVYRTDIKTIRDLRYSDRQWL
INLNNKIRSIVPGCYNYAVHPDELRILVHYQPSYYHFHIHIVNIKHPGLG
NSIAAGKAILLEDIIEMLNYLGPEGYMNKTITYAIGENHDLWKRGLEEEL
TKQLERDGIPKIPKIVNGFK*
```

**Supplementary Figure 4.** The 370 amino acid yeast DcpS as expressed with an amino terminal His tag. The HIT motif has been bolded.

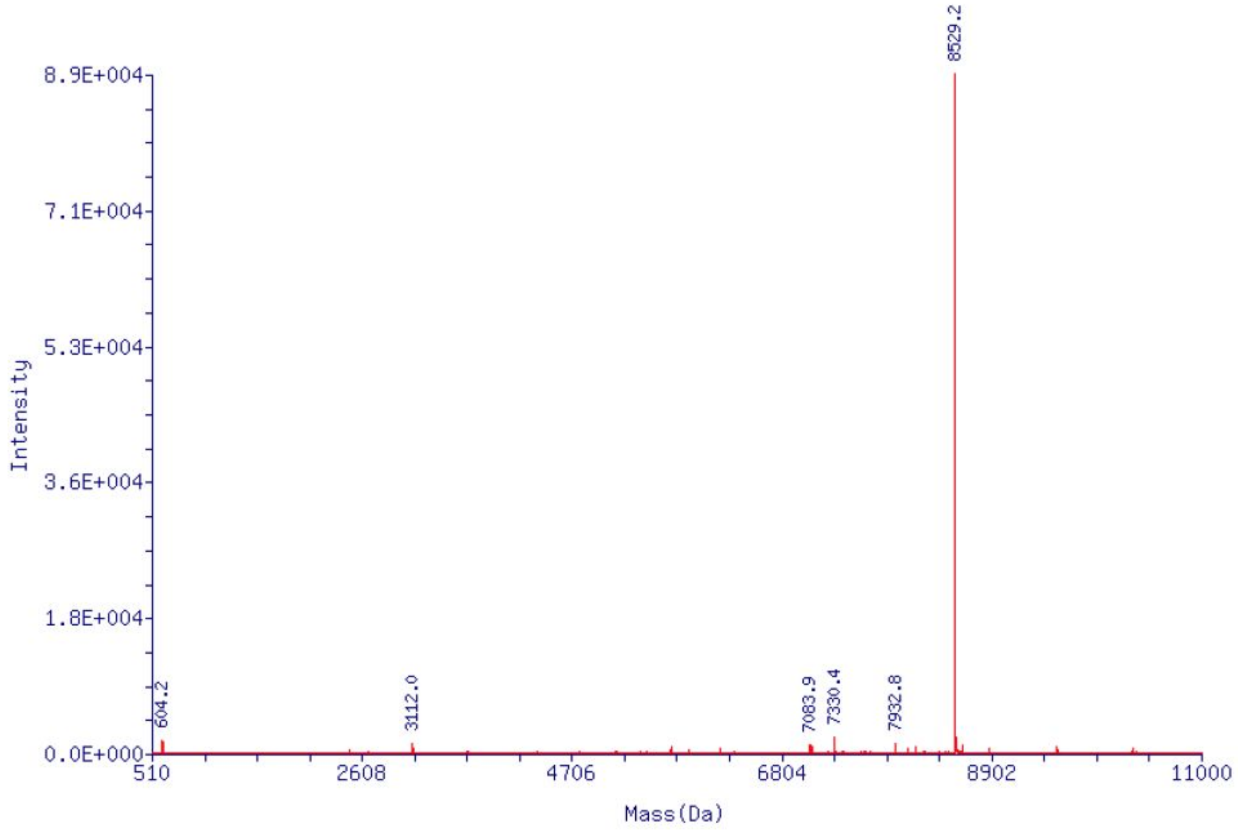


**Supplementary Figure 5.** Deconvoluted LTQ-XL mass spectrum of trimethylated $m^{2,2,7}$GpppG 25mer RNA obtained from the reaction of $m^7$GpppG 25mer RNA with hTGS. Target Mass (Da) calculated for $m^{2,2,7}$G*ppp*GUAGAACUUCGUCGAGUACGCUCAA ($C_{251}H_{315}N_{100}O_{185}P_{27}$): 8529.1. Observed Mass (Da): 8529.1. Mass Error: 0.1 Da (0.001%). Estimate Purity. 88.5%.

Full images for Figure 4b

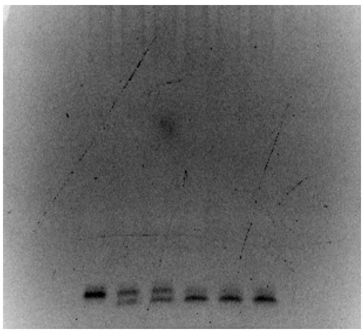

$m^7Gppp^6mA$

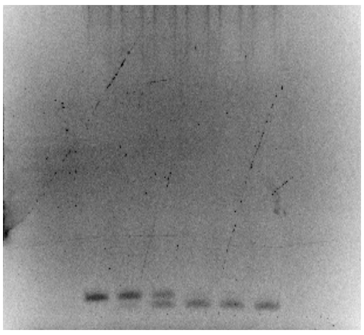

$m^7GpppG$

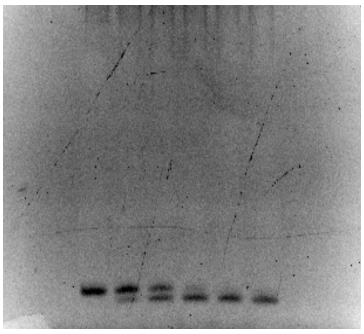

$m^7GpppGm$

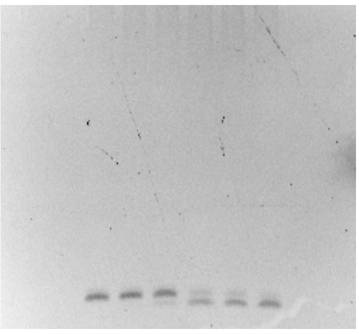

GpppG

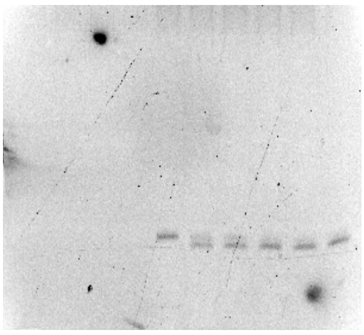

$m^7Gppp^6mAm$

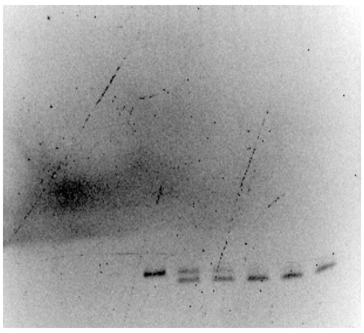

$m^7GpppA$
